# Single cell q-PCR derived expression profiles of identified sensory neurons

**DOI:** 10.1101/560672

**Authors:** Peter Adelman, Kyle Baumbauer, Robert Friedman, Mansi Shah, Margaret Wright, Erin Young, Michael P. Jankowski, Kathryn M. Albers, H. Richard Koerber

## Abstract

Sensory neurons are chemically and functionally heterogeneous and this heterogeneity has been examined extensively over the last several decades. These studies have employed a variety of different methodologies, including anatomical, electrophysiological and molecular approaches. Recent studies using next generation sequencing techniques have examined the transcriptome of single sensory neurons. Although, these reports have provided a wealth of exciting new information on the heterogeneity of sensory neurons, correlation with functional types is lacking. Here, we employed retrograde tracing of cutaneous and muscle afferents to examine the variety of mRNA expression profiles of individual, target-specific sensory neurons. In addition, we used an ex vivo skin/nerve/DRG/ spinal cord preparation to record and characterize the functional response properties of individual cutaneous sensory neurons that were then intracellularly labeled with fluorescent dyes, recovered from dissociated cultures and analyzed for gene expression. We found that by using single cell qPCR techniques and a limited set of genes, we can identify transcriptionally distinct groups. We have also used calcium imaging and single cell qPCR to determine the correlation between levels of mRNA expression and functional protein expression and how functional properties correlated with the different transcriptional groups. These studies show that although transcriptomics does map to functional types, within any one functional subgroup, there are highly variable patterns of gene expression. Thus, studies that rely on the expression pattern of one or a few genes as a stand in for physiological experiments, runs a high risk of data misinterpretation with respect to function.

**Significance statement:** Expression profiles of unidentified sensory neurons have been recently studied using RNASeq techniques. Here, we utilize a multifactorial approach to target identified cutaneous and muscle afferents to examine expression and functional levels of specific high priority candidate genes using *ex vivo* electrophysiology, Ca2+ imaging, and single cell qPCR. Using this methodology, we were able to identify specific groups of neurons with distinct functional properties that corresponded to unique transcriptional profiles. This represents the first attempt to relate neuronal phenotype with levels of gene expression in single identified afferents and highlights the importance of combining functional analysis with transcriptomics.

## Introduction

Understanding complex diseases of the nervous system, like chronic pain, requires a detailed analysis of the molecular components of the cells that contribute to pain. Within the peripheral nervous system, primary afferents are the principal transducers of environmental stimuli, and serve as the initial pathway by which noxious signals, in the form of action potentials, arrive in the central nervous system (CNS). How this information is interpreted and processed by cells within the CNS depends, in large part, on the functional properties of the sensory neuron responsible for transduction and transmission of this information. Primary afferents are a heterogeneous population of neurons comprised of cells with multiple neurochemically and functionally distinct identities that contribute to how they transduce environmental stimuli. There is a long history of attempts to link identified anatomic/neurochemical cellular characteristics to a cell’s functional properties, but progress in this domain has been hampered by this cellular complexity.

Concerted efforts have been made to determine the unique sets of biomarkers that correspond to nociceptive and non-nociceptive neurons. Traditionally, this has been attempted by examining various proteins expressed in cells using immunohistochemical methods. Technological advances, including next generation RNA sequencing, have expanded the definition of cellular identity, normalizing the use of transcriptional profiling and providing unparalleled insight into the transcriptional profiles within respective populations of nociceptive and non-nociceptive afferents. While these discoveries have provided a wealth of information, there still has not been a clear link made between transcriptional profile and functional properties (e.g., Usoskin et al. 2015, Zeisel et al., 2018). An additional complexity that has made this latter point challenging has been the limited use of transcriptional profiling in afferents that have been physiologically characterized within their native environments where their central and peripheral sites of termination remain intact. Examining afferent function and subsequent mRNA profile within the context of the peripheral tissue each cell innervates, is critical because previous work has shown that the tissue innervated influences the way in which nociceptors respond to inflammation and injury (Gold and Flake 2005; Malin et al. 2011).

Here, we utilized a multifaceted approach that employed retrograde tracing of cutaneous and muscle afferents to examine the variety of expression profiles of individual, target-specified sensory neurons. We characterized the expression of 28 high priority candidate genes known to contribute to the neuronal response to inflammation and injury using single cell real-time qPCR. Using this technique, we revealed significant diversity in patterns of expression between sensory neurons. We then utilized our *ex vivo* skin/nerve/DRG/spinal cord preparation to record and characterize the physiological response properties of individual cutaneous sensory neurons. Characterized neurons were collected and subjected to single cell qPCR analysis of transcriptional profile. Using this approach, we were able to identify collections of genes and expression levels that correlated with physiological response properties of each neuron. Our results represent the first time that transcriptional profile has been correlated with neuronal responses to quantitative naturalistic environmental stimulation, and provide additional insight into the genes that contribute to response properties of nociceptive afferents.

## Materials and Methods

### Animals

Experiments were conducted on adult (4-8 week) Swiss-Webster mice (Hilltop Farms or The Jackson Laboratory). Animals were group-housed with a 12-hour light-dark cycle and ad libitum access to food and water. All procedures were approved by the Institutional Animal Care and Use Committee at the University of Pittsburgh and used in accordance with AAALAC-approved practices.

### Retrograde labeling

Mice were anesthetized with isoflurane (2%) and an incision was made on the medial surface of the right thigh. The saphenous or femoral nerve was isolated with a minimum amount of soft tissue damage. Parafilm was placed behind the isolated nerve and the chosen dye (1% Alexa 488-Isolectin binding protein 4 [IB4], 1% Alexa 488-wheat germ agglutinin [WGA] or 0.5% cholera toxin subunit β [CTB]) was pressure injected into the appropriate nerve using a quartz pipette and a picospritzer. The wound was closed using silk sutures.

### Single cell survey design

Four animals per dye were back-labeled 24 hours before DRG collection. On the collection day, their L2 and L3 DRGs were removed and dissociated. Then 8-10 back-labeled cells were randomly selected and picked up from each animal and stored at −80ᵒC until processed further. Each cell’s RNA was then amplified and underwent qPCR until 6-8 viable samples (defined as having a GAPDH cycle threshold (Ct) of less than 25) were obtained for each animal.

### Cell dissociation and pickup

DRGs containing labeled cells were removed and dissociated as described previously (Malin et al., 2007). Briefly, DRGs were treated with papain (30 U) followed by collagenase CLS2 (10 U) /Dispase type II (7 U), centrifuged (1 minute at 1000 RPM), triturated in MEM, plated onto laminin-coated coverslips in 30mm diameter dishes and incubated at 37ᵒC for 45 minutes. Dishes were removed and flooded with collection buffer (140mM NaCl, 10mM Glucose, 10mM HEPES, 5mM KCl, 2mM CaCl_2_, 1mM MgCl_2_). Single, labeled cells were identified using fluorescence microscopy, picked up using glass capillaries (World Precision Instruments) held by a 4-axis micromanipulator under bright-field optics and transferred to tubes containing 3uL of lysis buffer (Epicentre, MessageBOOSTER kit). Cells were collected within 1 hour of removal from the incubator and within 4 hours of removal from the animals.

### Single Cell Amplification and qPCR

The RNA isolated from each cell was reverse transcribed and amplified using T7 linear amplification (Epicentre, MessageBOOSTER kit for cell lysate), run through RNA Cleaner & Concentrator-5 columns (Zymo Research), and analyzed using qPCR as described previously (Jankowski et al., 2009) using optimized primers and SsoAdvanced SYBR Green Master Mix (BIO-RAD). Threshold cycle time (Ct) values were for each well.

### Primer design and validation

Forward and reverse primer sequences were chosen for each gene within 500 bases of the 3’ poly A addition site. Primer sequences are available on request. Primer testing was done using cDNA generated using RNA from the whole DRG, as described in Jankowski et al., 2009. As described above for single cells,10 or 160pg aliquots of the RNA was amplified as described previously for single cells. For each primer set, serial dilutions of these aliquots were used to calculate primer efficiencies over the range of RNA concentrations observed in single cells as described in Pfaffl, 2004. Expression of specific genes was corrected for these primer efficiencies (Pfaffl, 2004) and expression level relative to GAPDH determined.

### Skin-nerve-DRG ex vivo preparation

The ex vivo preparation has been described previously (McIlwrath et al., 2007; Lawson et al., 2008). Briefly, mice were anesthetized with a mixture of ketamine (90mg/kg) and Xylazine (10mg/kg) mixture and perfused using 10-12ᵒC oxygenated (95% O_2_/5% CO_2_) ACSF (127mM NaCl, 26mM NaHCO_3_, 10mM D-glucose, 2.4mM CaCl_2_, 1.9mM KCl, 1.3mM MgSO_4_, 1.2mM KH_2_PO_4_). The spinal column and right hindlimb were removed and placed in a circulating bath of the same oxygenated ACSF. The shaved hairy skin of the hindlimb with connected saphenous nerve, L1-L5 DRGs, and the associated spinal cord were isolated and transferred to a second recording chamber with circulating oxygenated ACSF, where they were heated to 31ᵒC.

DRG neurons were impaled using quartz microelectrodes (>200 MOhm) containing 0.05% Alexa dye (555 or 488 nm) in 1M potassium acetate. Electrical search stimuli were delivered using a suction electrode on the saphenous nerve. Mechanical receptive fields were located by probing with Von Frey filaments or glass rod. If no mechanical receptive field was found, thermal fields were detected by applying hot (52ᵒC) or cold (0ᵒC) saline (0.9%) to the skin.

Once the receptive field was located, peripheral conduction velocity was calculated using spike latency and the distance between stimulation and recording electrodes. Controlled mechanical stimuli (square waves) were presented using a force-modulating mechanical stimulator (Aurora Scientific) with a 1mm diameter plastic foot. Thermal stimuli (rapid cooling to 4ᵒC or a 12s heat ramp from 31 to 52ᵒC) were presented using a 3mm Peltier element (Yale University Machine Shop). Cells were given 30 seconds to recover between stimuli. After electrophysiological characterization, cells were iontophoretically filled with Alexa dye. Only one cell per dye was injected per ganglion. Following the experiment, DRGs were removed, dissociated, and the labeled cells were collected as described above. Responses were analyzed offline using Spike2 software, Cambridge Electronic Design).

### Automated hierarchical clustering

Back-labeled and functionally characterized cells were clustered using the unweighted pair group method with averaging (UPGMA) using the expression information obtained from single cell PCR. Preprocessing for this data analysis consists of taking the ΔCt values and replacing the samples that failed to generate a value for a given gene with the detection limit for that gene. We then used MATLAB’s UPGMA implementation to agglomeratively cluster these neurons based on the Euclidean distance between their ΔCt values for each gene.

### Calcium imaging of dissociated neurons

Back-labeled cutaneous neurons were dissociated from L2 and L3 DRGs as described above and incubated with 2µM Fura2AM calcium indicator dye dissolved in Hank’s balanced salt solution HBSS for 30-60 minutes before recording. Coverslips were moved to an inverted fluorescence microscope (Olympus) where they were continuously superfused with HBSS using a tribarrel drug application device (Exfo Burleigh PCS-6000). A suitable field of view was located and ROIs were marked over cells identified with the white light transmittance.

Experimentation and shutter control was automated (Lambda DG-4/Lambda 10-B, Sutter Instruments) and two application protocols were used: 1 second 50µM αβmeATP, 1 second 50mM K^+^ and 1 second 1µM capsaicin, or 5 seconds 100µM 2-methylthioADP, 1 second 50mM K^+^ and 1 second 1µM capsaicin. Following imaging, a 3-axis micromanipulator with borosilicate glass electrodes was used to pick up cells into 3µL of lysis buffer (Epicentre, MessageBOOSTER), their ROI identifier was recorded, and the cells were stored at −80ᵒC until they could be amplified.

### Statistical analysis

Comparisons for levels of gene expression were performed using t-tests or Fisher’s exact test. Relationships between gene expression and Ca2+ influx were determined by Pearson’s correlation.

## Results

We recovered and an amplified the mRNA from a total of 247 sensory neurons from 64 mice. Of these, 120 were cutaneous afferents, 81 were back-labeled from the saphenous nerve (12 mice) and 39 were identified and intracellularly labeled in our *ex vivo* preparation (36 mice). An additional 127 cells were muscle afferent fibers back-labeled from the femoral nerve (16 mice).

### Validation of linear preamplification

To verify that pre-amplification was independent of both the specific RNA sequence being amplified and the starting amount of the specific mRNA, we pre-amplified 8 replicates of two different amounts of mRNA (10pg and 160pg) from Swiss-Webster lumbar whole DRG lysate. After pre-amplification, we pooled the replicate samples and proceeded through RNA column purification and qPCR, running twenty of our primer sets. The variable abundances of different mRNA templates in whole DRG lysate resulted in a wide distribution of Ct values, which allowed us to assess whether the amplification was dependent on the amount of starting transcript. As shown in Fig. 1, the 16-fold difference between starting amounts of mRNA resulted in amplification not significantly different from linear (Runs test, p=0.59) with a slope of 1.03 ± 0.05 indicating that the cDNA yield of amplification depended linearly on the starting mRNA amount. The 16-fold change in starting material would be predicted to impart a 4 Ct offset (2^4^ = 16), which was also supported by the observed 3.98 ± 1.186 Ct offset. Thus, we concluded that the preamplification of each mRNA is linear to the extent of our ability to detect it.

**Figure 1.**
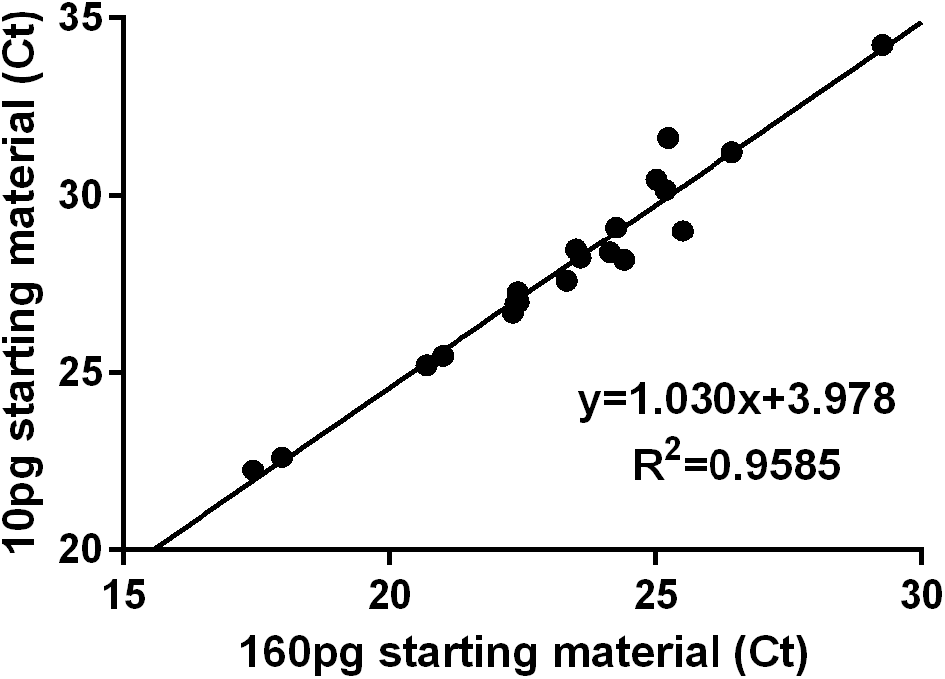
Preamplification linearity. Cycle thresholds obtained from qPCR of preamplified and pooled 10pg and 160pg starting material for different primer sets, corrected for primer efficiency.

### Clustering cutaneous afferent types by single cell qPCR

We identified 81 cutaneous neurons for single cell qPCR by back-labeling the saphenous nerve with commonly used tracers (IB4 n=14, WGA n=56, and CTβ n=11). Another 39 were collected and analyzed after being functionally characterized in our *ex vivo* skin-nerve-DRG preparation. This approach allowed the generation of a comprehensive distribution of a total of 120 cutaneous afferents, which were then clustered autonomously. Following the cluster analysis, the individual clusters were characterized by their expression profiles and each different group assigned a name. The emergent groups we identified were most similar to the classification scheme put forth by Usoskin et al. 2015 and Zeisel et al. 2018.

In our analysis, the root split in the cell dendrogram (Fig. 2) divides the identified cutaneous neurons into two groups based on the relative level of expression of the myelinated neuron marker neurofilament heavy chain (Nefh), a part of the NF200 complex. The low-Nefh expression cluster can be further subdivided into six populations of putative and characterized unmyelinated fibers, each with an identifying gene marker. For example, the green color-coded population expresses MAS-related G-protein coupled receptor, member D (*Mrgprd)*, lacks classic biomarkers for peptidergic neurons (e.g. Calcα (CGRP) and Tac1(Substance P)). This group also contains some of the highest levels of P2rX3, Asic2, and Gfrα2, as would be expected from the extensive literature on this population (Dong et al. 2001; Rau et al. 2009; Zylka et al. 2005).

**Figure 2.**
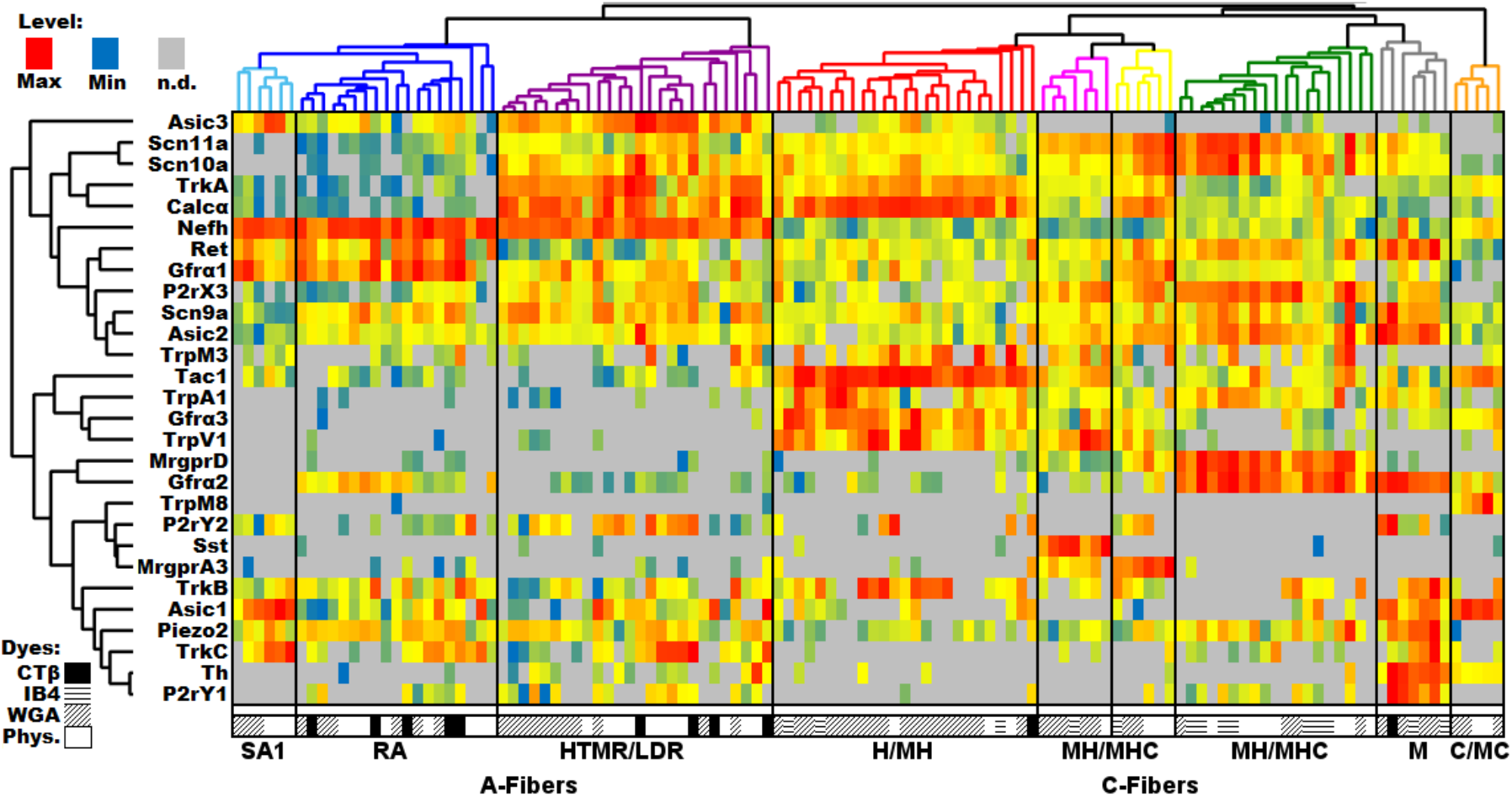
DRG profiling of identified cutaneous afferents using single cell PCR. Single cell qPCR results from identified cutaneous sensory neurons, with each column being a unique cell and rows representing genes. Heatmap temperatures for the histogram are scaled to expression, with red representing the maximum level, yellow representing 1/32 of the maximum level, blue representing the minimum level, and grey representing an undetectable level of transcript. Distributions were typically left-skewed, so scaling the heatmap in this way ensures the upper mode of the distribution will be colored red, orange, or yellow, while the lower tail is green or blue. Both cells and genes have been clustered using the same UPGMA algorithm. The symbols below each column indicates the source of each cell.

The pink-coded group of neurons is denoted by the high level of somatostatin (Sst) expression. Although the Sst group also expresses significant levels of Mas-related G-protein coupled receptor, member A3 (*Mrgpra3*), it can be distinguished from the yellow-coded cluster of neurons with higher levels of *Mrgpra3* expression, by its *Sst* expression and lower levels of *Trka*. Both clusters contain some level of *Mrgpra3* and *Trpv1* with low to undetectable levels of *Gfrα2*. The *Sst* population has been previously identified as being separate from the canonical “peptidergic” substance P (*Tac1*) population (Hökfelt et al. 1976), but the uniqueness of these cells was not fully appreciated until recently (Usoskin et al. 2015; Stantcheva et al. 2016). Similarly, *Mrgpra3* was identified as a population unique from the *Mrgprd* population that is positive for TRPV1, P2X3, CGRP, and c-RET protein expression, but not NF200 complex or substance P (Liu et al. 2009; Han et al. 2013). Although on average there is 11 times more *Mrgpra3* expressed in our A3 (yellow) subpopulation than in the Sst-containing group (t-test: p=0.011), *Mrgpra3* has been reported to co-localize with another marker for the Sst group, natriuretic peptide B (*Nppb*) (Mishra & Hoon 2013; Huang et al. 2018). Thus, there may be functional *Mrgpra3* expression in both populations. As the MrgprD, MrgprA3 and Sst groups exhibit prevalent labeling with IB4, they are likely to be the closest analogs for the population that was previously identified based on staining with the Isolectin B4.

The red-coded Tachkykinin-1-containing (Tac1) cluster is likely a large component of the canonical unmyelinated cutaneous peptidergic neurons, which also express high levels of *Calcα*, *TrkA*, *Gfrα3* and *Trpv1* (Skofitsch and Jacobowitz 1985; Hokfelt et al. 1975; Tominaga et al. 1998; Bennett et al. 2006). Histological methods of identifying peptidergic afferents tend to rely on the products of these five transcripts, which also map to functional differences (e.g Lawson et al. 2008), indicating that at least for this population, our unbiased methods correspond to previous functional and anatomical classification schemes.

The grey-coded tyrosine hydroxylase-containing (TH) cluster, included other known markers of this population such as *Gfrα2*, *Ret*, *P2ry1*, *Piezo2*, *Trkc* and *Asic1* (Li et al. 2011; Usoskin et al. 2015). The relatively low prevalence of this population is not surprising given that only 10-15% of the mouse DRG neurons express *TH*. This cutaneous population is poorly labeled by IB4 (Brumovsky et al. 2006) and identified C-LTMR fibers that have been shown to express TH, do not bind IB4 or express CGRP (Albers et al., 2006; Li et al., 2011).

The final group within the low-*Nefh* expressing cluster is the orange-coded group which is primarily distinguished by the expression of transient receptor potential cation channel, member M8 (*Trpm8*). In addition to *Trpm8*, these cells express relatively high levels of *Asic1*, *Gfrα3*, and *Tac1*. Notably, these fibers express a complement of sodium channels that is unique among unmyelinated sensory neurons. They lack detectable levels of Scn11a (Nav1.9) and have low to undetectable levels of Scn10a (Nav1.8), both of which are typically associated with small afferents and are expressed at high levels in cells expressing low levels of *Nefh*.

The high-*Nefh* expressing cluster is also clearly split into classical peptidergic and non-peptidergic groups. The purple-coded *Calcα* group has high levels of *Calcα*, *Trka*, and *Asic3*. The non-peptidergic myelinated fibers are divided into two groups. The dark blue-coded cluster has high levels of *Gfrα1*, *Ret* and *piezo2* whereas the light blue-coded cluster expressing high levels of *Asic1*, *Trkc* and *Asic3*. This latter group appears to closely match the NF3 group reported by Usoskin et al. (2015).

### Relationship of Ca2+ to relative transcript levels

Results from the single cutaneous neuron qPCR analysis reveal a wide range of transcript expression levels between cutaneous afferents, with several thousand-fold differences between maximum and minimum levels. Additionally, small but measurable transcript levels are occasionally expressed in neuron types not thought to contain the associated protein (e.g. *Calcα* in the non-peptidergic *MrgprD* population). These observations suggest the possibility that our technique may be detecting transcripts that are not translated to protein. To address this issue and examine the relationship between function and transcript levels, we used calcium imaging and single cell qPCR to compare agonist responses in cultured neurons to the associated receptors’ transcript levels for individual neurons (Fig.3). For these experiments, we chose *Trpv1*, *P2rx3*/*P2rx2*, and *P2ry1* as target transcripts because their respective receptors possess specific agonists and evidence suggests they are differentially expressed across the DRG cell subpopulations. For example, P2X3 and P2X2 are reported to be expressed in nonpeptidergic C-fibers (Guo et al. 1999; Petruska et al. 2000); *Trpv1* should be expressed largely in the peptidergic population (Michael & Priestley 1999) and *P2ry1* should be expressed in both large and small cells (Nakamura & Strittmatter 1996; Burnstock, 2007).

**Figure 3.**
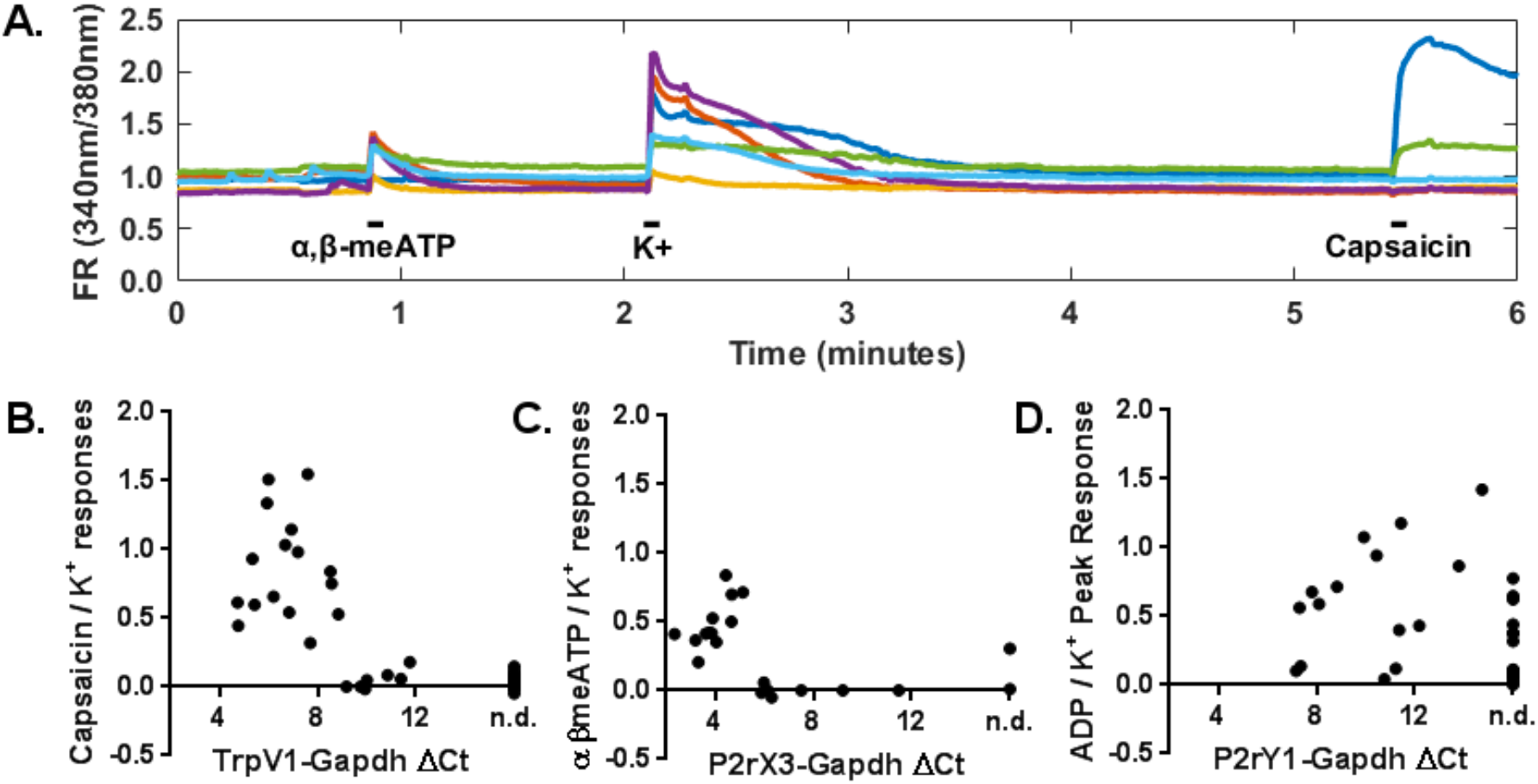
Calcium imaging reveals apparent thresholds of function for two ionotropic channels. (A) Example Fluorescence Ratio (FR) traces in response to 50µM α,β-meATP, 1M KCl, and 1µM Capsaicin (100mM ADP not shown). (B-D) Ratio of agonist to KCl responses versus relative receptor transcript expression for (B) *TrpV1*, (C) *P2rX3*, and (D) *P2rY1*.

In these experiments we examined transcript levels in cutaneous back-labeled cells that exhibited calcium transients evoked by α,β-meATP, capsaicin or ADP (Fig. 3A). Non-responsive cutaneous neurons, collected from the same cultures, were analyzed in parallel. Capsaicin responders and non-responders were clearly divided by their relative *Trpv1* transcript level (Fig. 3B). Neurons responsive to capsaicin always expressed detectable *Trpv1* transcript (16/16), while detection in non-responders was rare. Interestingly, several non-capsaicin sensitive cells were found to express low levels of *TrpV1* (7 of 23). Further analysis of this result showed a clear relationship between transcript levels and whether a cell would respond to capsaicin. Specifically, neurons that did not respond to capsaicin had *TrpV1* ΔCt levels over 9 (signifying lower levels of transcript expression), while every cell below 9 ΔCt responded to capsaicin (Fig. 3B).

Applying this threshold to findings from our single cell survey results demonstrated that in our *Tac1*, red-coded peptidergic group, where *TrpV1* expression levels ranged from ΔCt 4.57-10.39, (mean ΔCt = 6.5), that 15 of 25 cells should express functional levels. Likewise, for the *Mrgpra3* group (ΔCt 6.03-7.54, mean ΔCt = 7.01), 3 of 6 would be predicted to have functional TRPV1 and for the Sst group (ΔCt 4.58-7.05, mean ΔCt = 5.98), 5 of 7 would be predicted to contain functional TRPV1. Finally, in the MrgprD group, no cells (0/19 predicted functional) expressed *Trpv1* at functional levels. These results agrees with previously published immunohistochemistry, which primarily show *Trpv1* expression in CGRP^+^/NF200^−^ cells (Christianson et al. 2006; Guo et al. 1999; Michael and Priestley 1999). Among responders, however, there is no statistical correlation between ΔCt and absolute or relative response amplitude.

Similarly, *P2rx3* levels were related to cellular response to α,β-meATP **(3C)**. Only 1 of 9 neurons with a *P2rx3* ΔCt over 6 exhibited a significant α,β-meATP response, while all 11 neurons with a ΔCt below 6 was responsive to α,β-meATP (Fig. 3C). The responsive cell without *P2rX3* transcript had the highest recorded expression level of *P2rx2* (ΔCt = 6.70), another ATP receptor. *P2rx2* was only observed in 3 responsive neurons (ΔCt = 6.70, 10.82, 12.35) and the other 2 expressing *P2rx2*, were not responsive to α,β-meATP. Applying a 6 ΔCt threshold to findings from our single cell survey indicates that, within different groups, 16 of 19 Mrgprd, 5 of 6 Mrgpra3 and 6 of 7 Sst cells likely expressed functional levels of *P2rx3*. In the Tac1 group, only 2 of 22 cells expressed sufficient levels of *P2rx3*. Interestingly, 8 of 22 of myelinated cells expressed levels of *P2rx3* that would be predicted to be functional. This is not entirely unexpected, given that large or myelinated sensory neurons have been observed expressing *P2rx3* via *in situ* hybridization (Chen et al. 1995), immunohistochemistry (Bradbury et al. 1998), and function (Dowd et al. 1998; Hamilton et al. 2001). Although there was a clear threshold expression level for responding cells, there is no correlation between ΔCt and relative or absolute response amplitude.

Unlike the two ionotropic channels, relative expression levels of the metabotropic receptor *P2ry1* are poorly related to 2-methylthioADP responsiveness (Fig. 3D). Only 11 of 18 ADP responders expressed detectable levels of *P2ry1* (Average ΔCt 10.54), while 5 of 13 ADP nonresponders also expressed *P2ry1* (Average ΔCt 10.00). Neither the percent of cells with detectable expression (Fisher’s exact test: p = 0.29), nor the expression levels (t-test: p = 0.70) were significantly different between the samples.

### Distribution of functionally characterized cutaneous afferents within expression derived groups

Of the 120 identified cutaneous afferents depicted in Fig. 2, 39 were collected following characterization of their functional properties (Fig. 4). Of these, 39 cells were represented in all but one (TH) of the nine groups. We have separated out these fibers and report their expression profiles, conduction velocities and functional properties, including threshold data for different stimulus modalities (Fig. 4). This comparison shows that different types of characterized fibers consistently mapped to single cell clusters. For example, 3 characterized slowly adapting Type I mechanoreceptors (SAI) mapped together in the light blue-coded cluster. Likewise, myelinated RA fibers all mapped to the dark blue-coded cluster. The purple-coded cluster contains all the characterized myelinated nociceptors with high mechanical thresholds (HTMRs). This cluster also contains fibers that have low initial mechanical thresholds. Many of the latter fibers had initial thresholds of 5mN, the same as the majority of well-defined low threshold mechanoreceptors. This group of fibers exhibit a rapidly adapting response to low threshold stimulation, which becomes slowly adapting as the intensities increase. These fibers are also capable of encoding the intensity of the mechanical stimulus over a large range of intensities. In order to reflect the functional properties of these cells we are classifying them as having a large dynamic range (LDR) and have separated the two functional subdivisions of the purple-coded group by a dashed line. In addition, there was one characterized C-fiber that mapped to this group. It expressed very high levels of TH, but differed from the TH group by the lack of *Gfrα2* and *Ret* expression and high levels of *Calcα* and *Trka*.

**Fig. 4.**
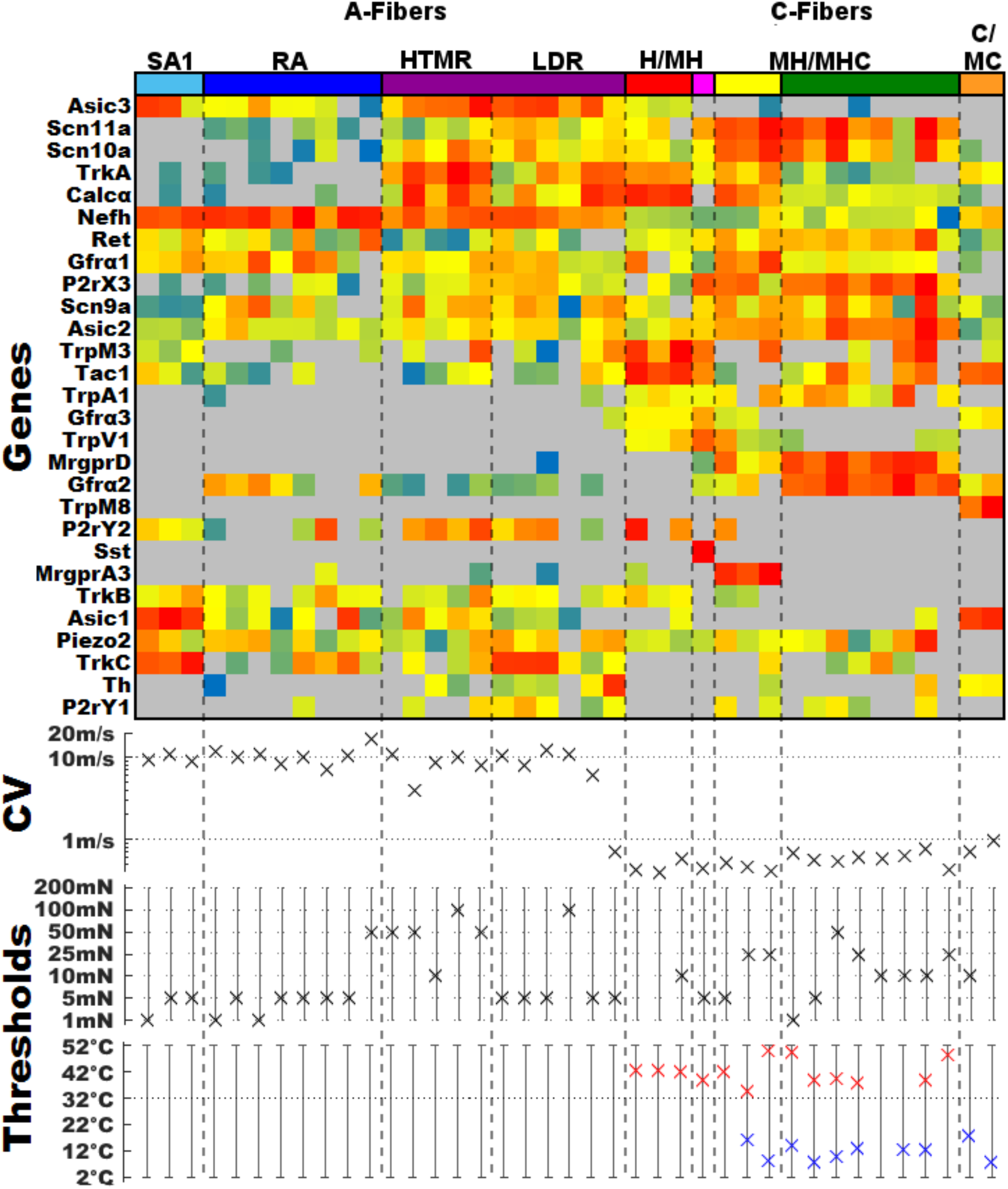
Expression profiles and the associated functional properties of characterized cutaneous neurons. *Top panel*: transcriptionally defined group color code and expression profiles of for individual characterized cells *Middle panel*: conduction velocity for each cell. *Bottom panel*: mechanical, heat and cold thresholds for each cell.

Within the C-fiber groups we found that the red-coded Tac1 group contained both mechanically insensitive, heat sensitive fibers (CH) and those that responded to both mechanical and heat (CMH). The three clusters corresponding to the classical non-peptidergic or IB4 binding population, Sst (purple), MrgprA3 (yellow) and MrgprD (green) contained cells that were classified as C-mechanical fibers (CM; no response to heat or cold), CMH (mechanical and heat, but not cold) and CMHC (C-mechanical, heat and cold). Furthermore, when comparing the response of the heat sensitive cells, those in the Tac1/Trpv1 group had higher peak instantaneous firing rates (range 50.8 – 56.4 Hz) compared to other heat sensitive fibers (range = 1.65 – 14.43 Hz) in response to heat stimulation, as has been reported previously when comparing the CH population to CPM population (Lawson et al. 2008)

The two characterized cells in the orange-coded Trpm8 group were both cold sensitive, and displayed higher peak instantaneous firing rates in response to cold (29.4 and 158 Hz) relative to the other 8 cold sensitive fibers (range = 0.65 – 5.14 Hz) which mapped to the MrgprD and MrgprA3 clusters. This agrees with our recent finding that *Trpm8* expressing cold sensitive fibers have similarly high instantaneous frequencies (Jankowski et al. 2017; also see Zimmermann et al. 2009; Campero et al. 2009). This unique property may be due in part to the cluster’s lack of the tetrodotoxin (TTX) insensitive *Scn10a* and *Scn11a* (Nav1.8 and 1.9, respectively).

### Clustering muscle afferent types by single cell qPCR

To compare results from cutaneous afferent profiling, we collected 113 identified muscle afferents following back-labeling from the femoral nerve using WGA (n=93) and CTβ (n=20). We used the same clustering procedure to autonomously place the individual cells into 7 defined groups (Fig. 5). In the construction of the heat map for muscle afferents we used the same ordering of genes to allow for easier comparison with the cutaneous neurons. This cluster analysis produced fewer groups than determined for cutaneous afferents. The expression patterns of 28 genes analyzed in individual muscle afferents showed a greater heterogeneity within some of these muscle afferent clusters when compared to their cutaneous counterparts. This is most obvious in the large red-coded peptidergic group that is denoted by the expression of high levels of *Tac1* and/or *Calcα*. This group appears to contain several subgroups. Most obvious is the subgroup expressing relatively high levels of *Nefh*, suggesting that they may be myelinated fibers, possibly Group III. This group also appears to be distinct from the other clusters by its moderate expression of *ASIC3* but low levels of *TRPV1*.

**Fig. 5.**
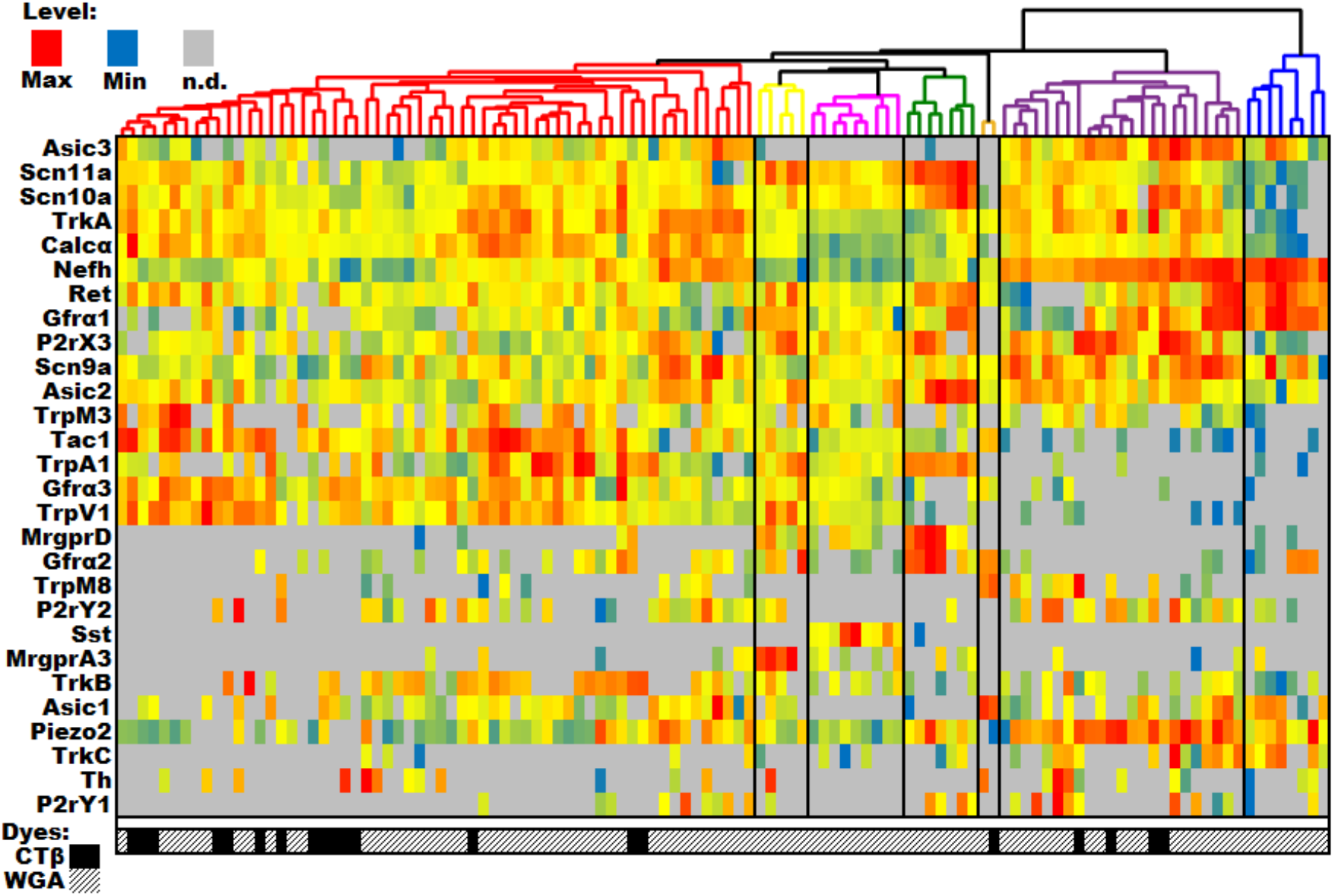
DRG profiling of identified cutaneous afferents using single cell PCR. Single cell qPCR results from identified muscle sensory neurons, with each column being a unique cell and rows representing genes. Heatmap temperatures for the histogram are scaled to expression, with red representing the maximum level, yellow representing 1/32 of the maximum level, blue representing the minimum level, and grey representing an undetectable level of transcript. Distributions were typically left-skewed, so scaling the heatmap in this way ensures the upper mode of the distribution will be colored red, orange, or yellow, while the lower tail is green or blue. The genes have been clustered using the UPGMA algorithm. The genes are listed in the same order as the cutaneous neurons in Fig. 2 for ease of comparison. The symbols below each column indicates the source of each cell.

Three other potential subgroups within the red-coded group that appear to be comprised of unmyelinated Group IV muscle afferents. Two of these more peptidergic groups have much in common with their cutaneous counterparts, co-expressing high levels of *Trpv1*, *Trpa1* and *Gfrα3*. The primary difference between the two groups appears to be differences in *TrkB* transcript levels and possibly *ASIC3* expression. The third group of putative group IV fibers express lower, or undetectable levels, of *Tac1*, *Calcα*, *Asic3* and *Trpa1*. Interestingly many of these cells were obtained by CTB back-labeling that is typically used to target myelinated, somatic afferents. However, it is also effective at labeling visceral afferents (e.g. Christianson et al 2006; Fasanella et al 2008).

There were also several muscle afferent groups that exhibited unexpected similarities with cutaneous C-fibers groups. Most notably we observed MrgprD (green) and MrgprA3 (yellow) expressing cells that were thought to be restricted to the cutaneous population. The relative expression of all 28 genes analyzed here were consistent across cutaneous and muscle afferents within these groups. *Sst* expressing neurons (magenta) were also present in both populations. However, in this case the co-expression of other genes differed somewhat from the cutaneous group, most notably due to the lack of *Mrgpra3* expression. In addition, the orange TrpM8 group appears to be similar to the cutaneous counterpart including the lack of the tetrodotoxin (TTX) insensitive sodium channels *Scn10a* and *Scn11a* (Nav1.8 and 1.9, respectively) and the expression of *Tac1, Asic1* and *Gfrα2*. There were many other *Trpm8* expressing muscle afferents that mapped to different groups including putative myelinated fibers in the magenta group. This suggests that unlike the cutaneous population, *Trpm8* expressing muscle afferents represent more heterogeneous groups of afferent fibers.

One of the cutaneous C-fiber groups missing from muscle group IV fibers was the TH group. Interestingly, while the percentage of cells expressing high levels of *Th* was similar between the cutaneous and other populations of muscle afferent, the co-expression of other sampled genes differed in the *TH*-expressing muscle fibers causing them to be split into several of the other clusters.

The three remaining groups appear to be representative of myelinated fibers due to the high levels of *Nefh* expression. The blue-coded group has expression profiles indicative of proprioceptors including high levels of *Nefh* expression and little or no Scn11a or Scn10a. These cells also uniformly expressed high levels of *Ret* and *Gfrα1*. For the most part, the purple-coded group, has similar expression levels for many of the same genes including high levels of *Nefh*, *Asic2*, *Asic3* and *Piezo2*. This suggests this group represents some type of myelinated mechanoreceptors and possibly some that are chemoreceptors given the expression of *Asic3*, *P2rx3*, *P2ry1* and *P2ry2* in many of these afferents.

## DISCUSSION

### Clustering of identified single cutaneous sensory neurons

Here we have shown the ability to use a linear amplification protocol and qPCR methodology to assign cells to functional transcriptional groups using the relative expression of 28 genes. Our automated clustering algorithm isolated nine clusters of cutaneous sensory neurons. While previous RNASeq studies of single DRG neurons have used afferents with unspecified target tissues (Chiu et al. 2014; Usoskin et al. 2015; Li et al. 2015; Zeisel et al., 2018), we have identified eight transcriptionally defined classes of cutaneous afferents that closely resemble previously identified groups (Chiu et al. 2014; Li et al. 2015; Usoskin et al. 2015). However, a recent deep sequencing study (Zeisel et al 2018) has reported several additional groupings of DRG neurons. The clusters we have identified here will be discussed with reference to this study. For comparison we have mapped the 7-8 highest expressed genes in our individual clusters to the expression data available at (http://mousebrain.org/).

Using this approach our SA1 group maps to the PSNF2 group reported by Zeisel et al., 2018, whereas the LTMR-RA group corresponds to their PSNF1 group. Our LDR group matches the PSNF3 group and the myelinated HTMR cluster the PSPEP1 group. Among the C-fibers, our Tac1 cluster is analogous to their PSPEP3 group. The MrgprA3 cluster is matches the PSNP4 group, whereas our SST cluster also closely corresponds to their PSNP6 and the Th cluster is representative of their PSNP1 group. While they identified 3 TRPM8 groups, our cutaneous cluster most closely resembles their PSPEP8 group. Finally, our MrgprD cluster closely matched both the PSNP2 and PSNP3 groupings.

### Relationship between relative expression level and function

We have been able to demonstrate a strong relationship between the relative level of gene expression and function for two receptor/ion channels in individual cutaneous sensory neurons. Although we did not detect a statistical correlation between gene expression and magnitude of Ca^2+^ transients, there was well-defined level of gene expression where Ca^2+^ influx was observed. We also found that there was no relationship between the relative expression level of *P2ry1* in cells that responded to ADP. In fact, ADP application evoked significant calcium transients in 8 cells that did not contain measurable levels of *P2ry1*, suggesting that *P2ry1* mRNA may not be maintained at a constant level and is transcribed on demand or post-transcriptionally modified. Alternatively, there may be another ADP receptor.

### Functionally characterized cutaneous sensory neurons

Segregation of functionally characterized cutaneous sensory neurons in appropriate individual clusters adds further validation to these results. Within the A-fiber population, the Aß SAI and RA fibers exhibit narrow somal action potentials and separate into two distinct groups. The remaining myelinated fibers map to the purple-coded cluster where all the cells exhibited broad somal action potentials that contain myelinated peptidergic high threshold mechanoreceptors (HTMRs) and afferents functionally characterized as fast conducting myelinated fibers with low initial mechanical thresholds that also encode stimulus intensities into the noxious range (LDRs). Of the 6 neurons in the LDR group, 3 were found to express high levels of *Trkc*. These afferents may be analogous to TrkC positive fibers that form circumferential endings around hair follicles, as they respond to graded mechanical stimuli (Bai et al., 2015; see figure S5).

The other 2 myelinated fibers in this group express relatively high levels *Trka* and *Calcα*. It has recently been reported that there are two types of fibers that make circumferential endings around high follicles; one that contains TrkC and the other CGRP. Interestingly, both subtypes apparently responded most vigorously to hair tugging (Bai et al., 2015; Ghitani et al., 2017). While Bai et al., (2015) considered the TrkC expressing group to be low threshold mechanoreceptors based on their response to light stroking of the hairy skin, Ghitani et al., (2017) describe the *Calcα* expressing neurons as nociceptors due to the vigorous response to tugging on the hairs.

For the C-fiber clusters, the Tac1 cluster contained 2 CH fibers and 1 CMH fiber. The Sst, MrgprA3 and MrgprD clusters were similar in that all the cells were mechanically sensitive containing CM, CMH, CMC and CMHC fibers. This matches previous studies characterizing cutaneous fibers expressing *mrgprd* and *mrgpra3* (Rau et al 2006; Han et al 2013). Finally, the TrpM8 cluster contains cold sensitive cells. Fibers in his group have several characteristic separating them from others C-fibers. For example, cooling sensitive TrpM8 fibers express little or no *Nav 1.8* or *1.9*, suggesting that this population may have a complement of voltage gated sodium channels that is more similar to myelinated fibers. The lack of two major TTX resistant sodium channels also raises the possibility that these cells are the TTX sensitive small cold fiber population (Sarria et al. 2012). In addition, this difference in their voltage gated ion channel compliment could potentially underlie the high instantaneous frequency that these fibers display in response to cooling (Jankowski et al., 2017).

Another intriguing finding in the TrpM8 group is the expression of *Tac1*. It is present without significant expression of other classical markers of peptidergic fibers such as *Calcα*, *Gfrα3* or *Trka* (Wang et al. 2013; Dhaka et al. 2008). Earlier genetic studies on the distribution of TrpM8, were inconsistent with one group showing significant overlap with CGRP and substance P (Takashima et al., 2007), while the other reported no significant overlap with either CGRP or substance P (Dhaka et al., 2008). Interestingly, in the recent study by Zeisel et al (2018), Tac1 was co-expressed with TRPM8 in all 3 groupings of TRPM8 fibers. Intriguingly, the presence of *Tac1* in these fibers could provide insight into the mechanism of cold allodynia following injury. Release of substance P in lamina I could positively modulate activity in NK1R positive lamina I projection neurons. This finding also suggests the possibility that these cooling fibers could play a role in the thermal grill illusion. For example, Substance P release from these terminals in lamina I could increase sensitivity of adjacent NK1R containing neurons to relatively innocuous thermal inputs.

### Clustering of single identified muscle sensory neurons

Studies by Usoskin (2015) and Chiu (2014) have provided information on the molecular profiles of putative muscle afferent populations. However, these profiles were defined based on their expression of parvalbumin, a marker of proprioceptor afferents. No studies to date have defined the expression patterns of group III and IV fibers. Although our results for identified muscle afferents does not clearly define these conduction velocity groups, our data does provide meaningful information on putative myelinated and unmyelinated muscle afferent subpopulations.

Many of the high *Nefh* expressing muscle afferents also express high levels of *Asic3*. This channel is a known proton sensing channel but has also been linked to modulation of mechanosensation (Sluka et al 2003), nociceptive processing (Chen et al 2014; Walder et al 2010; Ross et al 2016; Naves and McCLeskey 2005) and modulation of cardiovascular reflexes (Tsuchimochi et al 2011). These cells also appear to contain the mechanosensitive receptor, piezo2 (Coste et al 2010), which may contribute to a mechano-sensory function of these particular myelinated afferents. Whether these are of the proprioceptor or group III mechanoreceptor subtype remains to be elucidated.

Of the low Nefh, putative unmyelinated muscle afferents, several groups were detected. The non-peptidergic (low *Tac1/Calcα*), putative group IV afferents were *Asic3* negative but could contain *TrpV1* and *P2rX3*. Both of these latter receptors have been shown to play roles in cardiovascular reflex regulation (Smith et al 2010; Li et al 2014; McCord et al 1985) and muscle pain (Saloman et al 2013; Shinoda et al 2008) and appear to be contained within a population of muscle afferents with responsiveness to muscle metabolites such as lactate, ATP and protons (Light et al 2008, Jankowski et al 2013; Ross et al 2016). These cells have been found to be of the putative metabo-nociceptor and innocuous metaboreceptor subtypes (Jankowski et al 2013; Ross et al 2014). While *Asic3* expression has been linked to each of these processes (Chen et al 2014; Walder et al 2010; Ross et al 2016; Naves and McCleskey 2005; Tsuchimochi et al 2011), its expression does not necessarily overlap with TRPV1 or P2Xr3 (Jankowski et al 2013). Our current data does show a population of unmyelinated muscle afferents that are peptidergic and express *Trpv1*, *Asic3* and *P2rx3*. It is possible that these cells more likely fall within the nociceptor muscle afferent subtype that may be able to respond to multiple stimulus modalities (Jankowski et al 2013). *P2ry1* did not appear to have robust expression in many muscle afferent subtypes, but this G-protein coupled receptor has been shown to regulate a variety of muscle afferent responses, including modulation of nociception, mechanical and chemical responses, and cardiovascular reflexes (Queme et al 2016).

The results presented here from single cell qPCR on identified and functionally characterized cells demonstrates the utility of this approach to unlock the potential of the vast amount of information contained in the recent next-gen RNAseq studies and those to come.

## Acknowledgements

Funding was provided by NINDS NS023725 (HRK) NIAMS AR069951 (KAA).

